# DNA Damage Induced Chromosomal Instability. Computational modeling view

**DOI:** 10.1101/218826

**Authors:** Y.A. Eidelman, S.V. Slanina, V.S. Pyatenko, S.G. Andreev

## Abstract

The origin of dose-response curves for radiation-induced chromosomal instability (CI) is studied using the mechanistic CI model. The model takes into account DNA damage generation and repair in the progeny of irradiated cells, cell passage through mitotic cycle, and intercellular signaling. It is shown that the dose-response curves are closely related to the dynamic curves. The principles underlying this relationship are analyzed.

## INTRODUCTION

Chromosomal instability (CI) is defined as the increased frequency of chromosomal rearrangements (chromosomal aberrations) in offspring of irradiated cells [1]. As a theoretical basis, the theory of targets is not applicable here [2]. The first attempts at computer simulation of CI brought about promising results [3,4], but led to a multitude of questions. Some of these issues are discussed in this paper. The influence of various parameters and factors on the main characteristics of gamma-induced dose-response and dynamic curves are taken into account and analyzed here. As a criteria of delayed effects, or CI, the delayed dicentrics and chromatid aberrations are modeled as an ends-point of radiation-induced CI. Compared to [4], an assessment is made of the influence of signaling factors that cause DNA damage as a result of interaction between cells and are not reducible to primary lesions induced by radiation. The sensitivity of dose-response curves for dicentrics to the variation of the parameters of the CI model is analysed.

## MODEL

The mechanistic model of CI following ionizing radiation exposure incorporates DNA / chromosome damage interaction pathways determining outcomes of all factors involved in genome destabilization. The main modelling technique described previously [4] is used here with some modifications. The main points of the model are as follows.

DNA double-strand breaks (DSBs) can be formed in both irradiated cells and their progeny in the G_1_ phase, as well as in the S phase. The DSB formation in the G_2_ phase upon irradiation is neglected. For most cell types, when the asynchronous population is irradiated, the fraction of G_2_ cells is small, 10-15%. In addition, irradiation of G_2_-phase cells does not lead to the formation of dicentrics.

In addition to DSBs, DNA single-strand breaks (SSBs) and oxidative base damage (BD), as well as complex lesions (SSB + BD) are also induced. They are repaired by BER pathway. Unrepaired SSBs + BDs alone do not lead to the formation of aberrations, but can turn into DSBs either due to nuclease attack of opposite DNA chain in G_1_ or during replication.

The DSBs can be repaired by NHEJ or HR pathways, misrepaired with the formation of aberration of chromosome or chromatid type depending on the phase of the cell cycle, or lose their reactivity, forming a blunt-end aberration, or fragment. In the progeny of irradiated cells, as in [4], we consider the formation of DSBs *de novo* in the S phase, where predominantly chromatid-type aberrations are formed, of which sister chromatid exchanges of the "isochromatid deletion" type, or "chromatid dicentrics" are of primary interest.

When the cell passes into mitosis, the fate of CAs depends on their types. Chromosomal and chromatid fragments, having passed into mitosis, either are transmitted into one of the daughter cells, or are lost. A chromatid dicentric in mitosis forms an anaphase bridge. Since both kinetochores in this aberration belong to the same chromatid, it cannot segregate normally. The mitotic spindle pulls it to opposite poles. An anaphase bridge either leads to cell death, or breaks (part of the so called “BFB cycle”), and each of the daughter cells gets a centric chromosomal fragment with a sticky end.

The chromosomal dicentric, in contrast to the chromatid dicentric, has four kinetochores, two on each chromatid, and therefore can either segregate normally, or form a double anaphase bridge. With normal segregation, each of the daughter cells receives a dicentric, and with the formation of a double bridge, the same outcomes are possible as in the case of a single bridge: the death and bridge breakage as part of the BFB cycle.

As a result of breakage of anaphase bridges, in G_1_ centric fragments with sticky ends appear: single or double, depending on the type of the bridge that is broken. The reactive ends can interact with each other as well as with the DSBs generated in the S phase to form dicentrics and other types of aberrations. The cycle of formation, breakage and fusion of anaphase bridges is called BFB (Breakage and Fusion of Bridges).

DSBs of non-radiation nature in the progeny are formed through three channels, N_dsb_ = N_1_ + N_2_ + N_3_. N_1_ is an autonomous channel, i.e. these DSBs are formed in each cell regardless of external factors (intercellular signals). N_2_ is a non-autonomous channel, determined by intercellular interactions through gap junctions. It depends on the density of the cells, both having and not having DNA damage. N_3_ is also a non-autonomous channel, it is determined by intercellular interactions through soluble factors in the medium. DSBs of all three types are considered structurally indistinguishable from each other; accordingly, all of them are subject to the same processes with the same rates.

The cell passage through cell cycle is taken into account in the same way as in [4].

## RESULTS

We studied impact of the parameters of spontaneous DNA double-strand break (DSB) generation in the progeny of irradiated cells and the parameters of the breakage-fusion of the anaphase bridge (BFB cycle) on the shape of the dynamic and dose curves of radiation-induced CI.

Figs 1 and 2 demonstrate that, depending on the combination of parameters, three main types of dynamic curves can be distinguished: with the plateau (Fig.1c) and without the plateau (Fig.1d,e, Fig.2a), which in turn are divided into two subtypes: leading to the plateau by dose (Fig.1f, Fig.2b) and not leading (Fig.1g,h). All curves are observed for the rate of DSB generation independent of the dose in the range of medium and large doses (Fig.1b).

**Fig.1.**
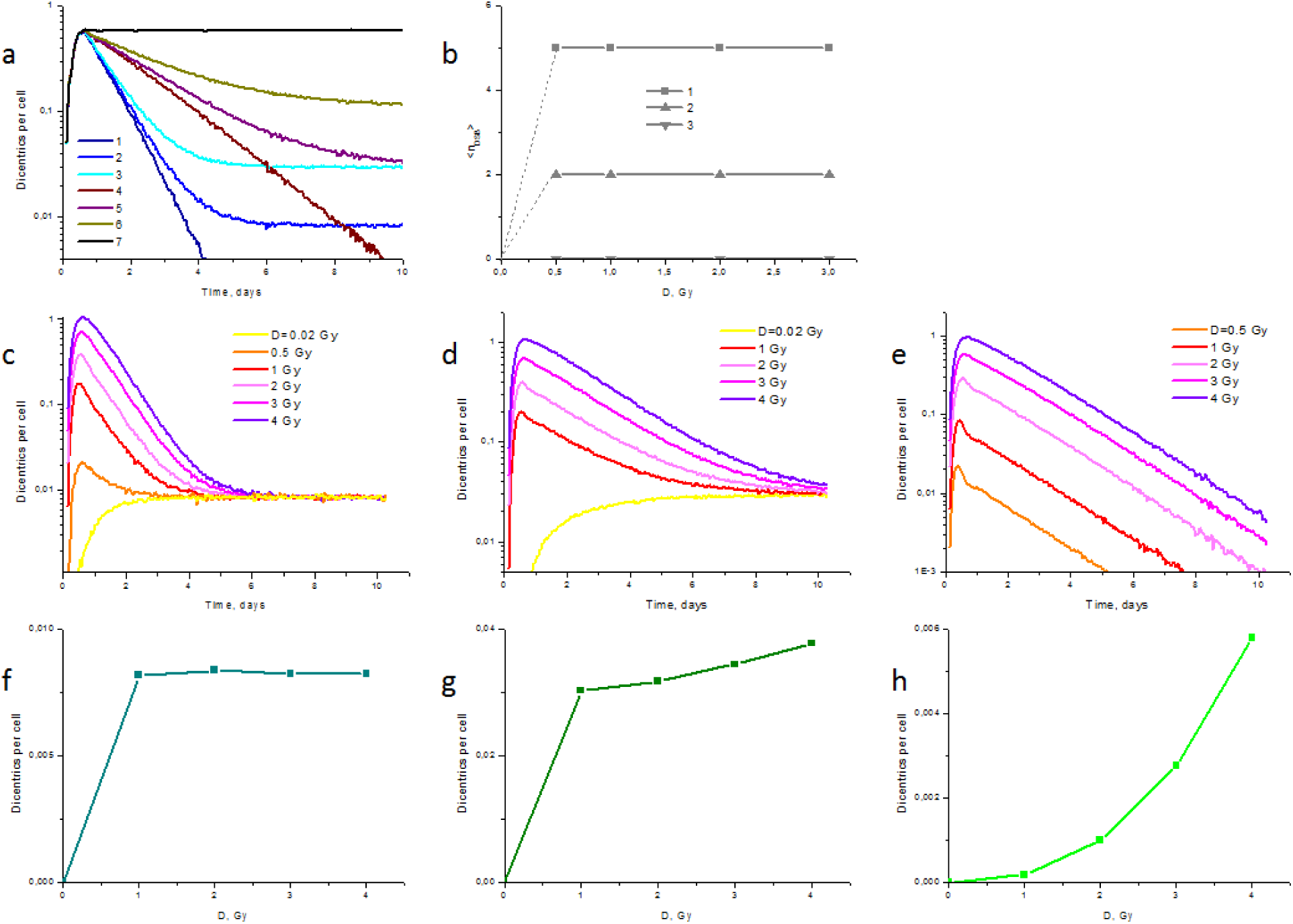
Impact of the DSB generation parameters and BFB on the shape of kinetic and dose curves of CI. (a): dynamic curves following 3 Gy irradiation for different probabilities of anaphase bridge breakage and different levels of DSB generation in the S phase. 1-3 - probability of bridge breakage *p*=0.6; 4-6 -*p*=0.3; 7 - *p*=1.0. 1, 4, 7 – mean number of DSBs <*n*>=0, i.e. no spontaneous generation; 2.5 - <*n*>=2 per S phase; 3.6 - <*n*>=5 per S phase. (b): dose dependencies of DSB generation in the S phase. 1 - corresponding to curves 3 and 6 on the panel (a); 2 - corresponding to curves 2 and 5 on the panel (a); 3 - corresponding to the curves 1, 4 and 7 on the panel (a). (c) - (e): dynamic curves for different doses with parameters corresponding to curves 2, 5 and 4 on the panel (a), respectively. (f) - (h): dose curves at t = 10 days, corresponding to the dynamic curves on the panels (c) - (e), respectively.

**Fig.2.**
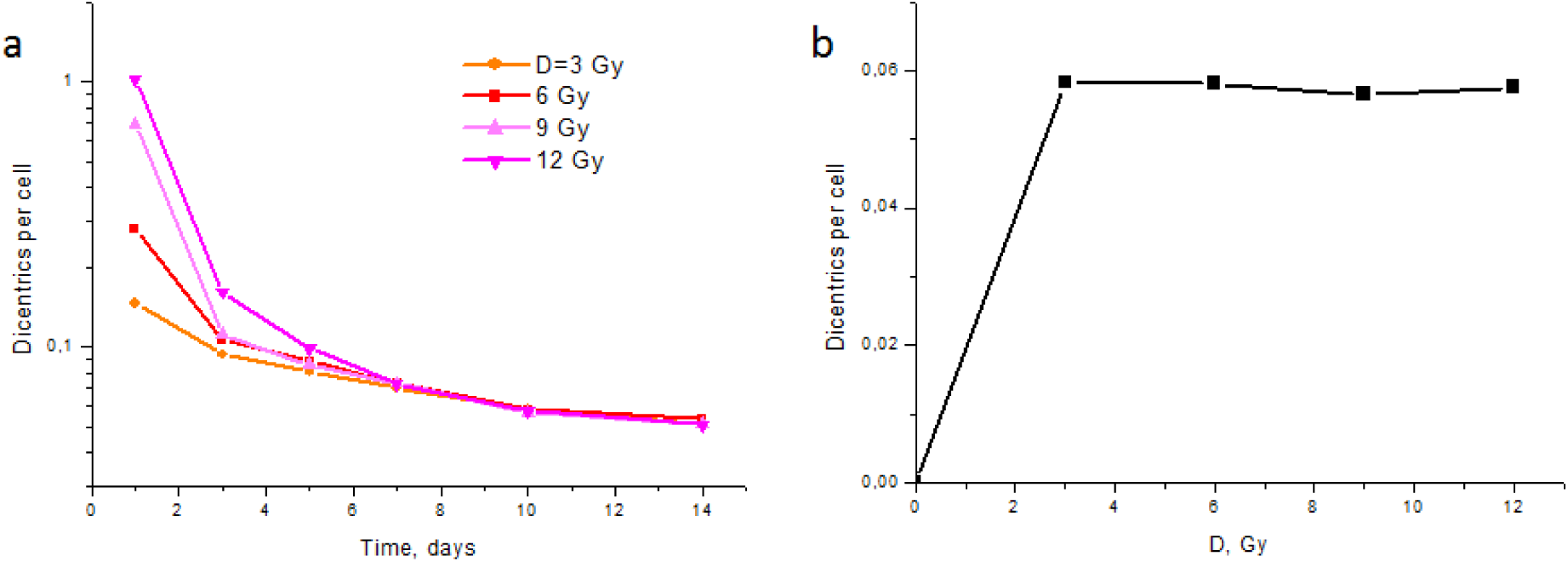
Plateau on the dose curve may not be accompanied by a plateau on the dynamic curve. (a): dynamic curves for different doses. (b): the dose curve at t = 10 days.

Next, we analyzed the sensitivity of the result, the dose dependence of the type seen in Fig.1f, to the pathway of spontaneous DSB generation: autonomous, i.e. independent of intercellular signaling, and the non-autonomous pathway. In the illustrative calculations presented in Figs 1 and 2 only autonomous DSB generation appears, since it does not depend on time, unlike intercellular signaling. Next we will assess how non-autonomous generation of DSBs affects the dynamic and dose curves.

The dynamic curves presented in Fig.3a are obtained for the same level of autonomous DSB generation, but the varying level of non-autonomous generation, caused by soluble factors in the medium. With the increase of non-autonomous generation, the kinetic curves go higher, and, in addition, the plateau is replaced by a wavy curve. The latter is due to the contribution of soluble factors. In the calculations of proliferation the cells are replated every two days (typical experimental conditions of cultivation). Immediately after the replating, the contribution of soluble factors to the DSB generation is absent, then increases until the next replating. Given the stochasticity of transitions between the cell cycle phases (i.e., the time when the generated DSB is manifested as a dicentric in mitosis, is different for each individual cell), this results in a wave-like dynamic curve of dicentrics.

In this case, the dose curves vary only in height (the larger the contribution of the non-autonomous DSB generation, the higher they are) but not in shape: the plateau observed for a purely autonomous mechanism is preserved even when the autonomous mechanism is switched on, Fig.3b. A similar conclusion is valid for the second non-autonomous channel, the formation of DSBs due to intercellular signaling via gap junctions, which can be seen in Fig.4.

**Fig.3.**
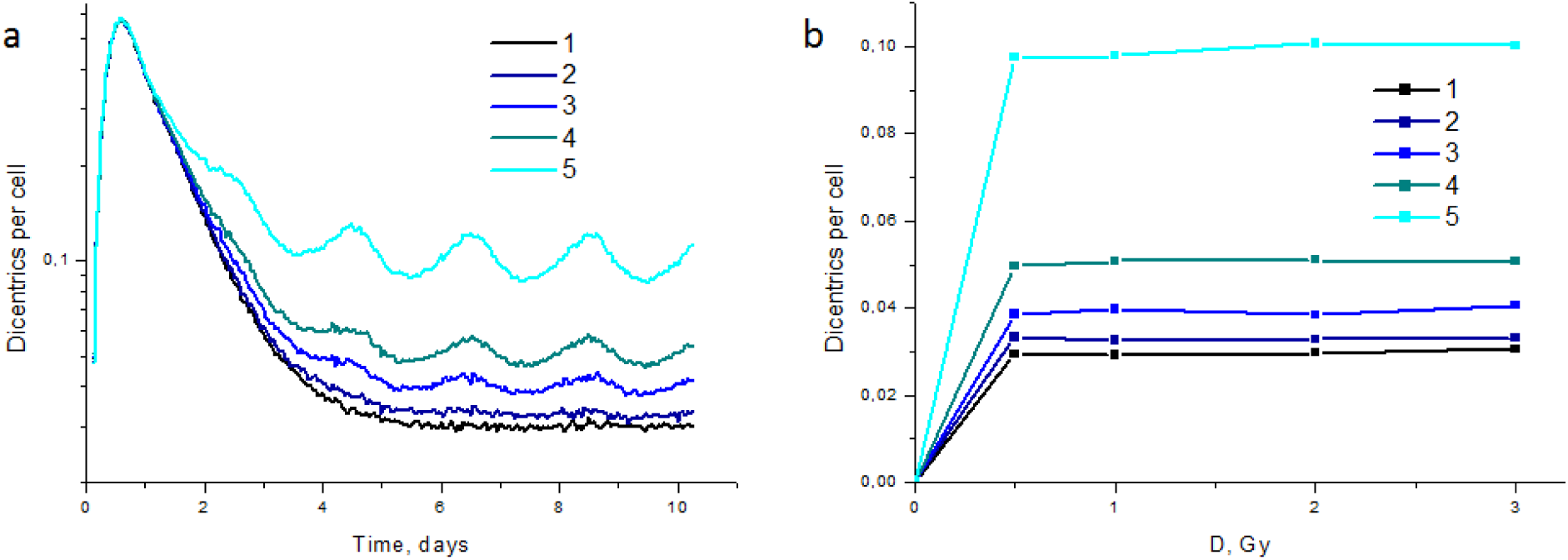
Non-autonomous DSB generation due to soluble factors affects the dynamic curves of CI, but does not change the shape of the dose curves. (a): dynamic curves at a dose of 3 Gy for different contributions of autonomous and non-autonomous pathways of DSB generation. The autonomous level in all cases <n> = 5 per S phase. The level of non-autonomous DSB generation due to soluble factors released by cells into the medium is varied. 1 - non-autonomous generation is absent (corresponds to curve 3 in Fig.1a). 2 - <n_SF_> = 0.43 per S phase; 3 - <n_SF_> = 1.46; 4 - <n_SF_> = 4.17; 5 - <n_SF_> = 8.74. (b): dose dependencies at t = 8 days. The designations of curves are the same as in (a).

**Fig.4.**
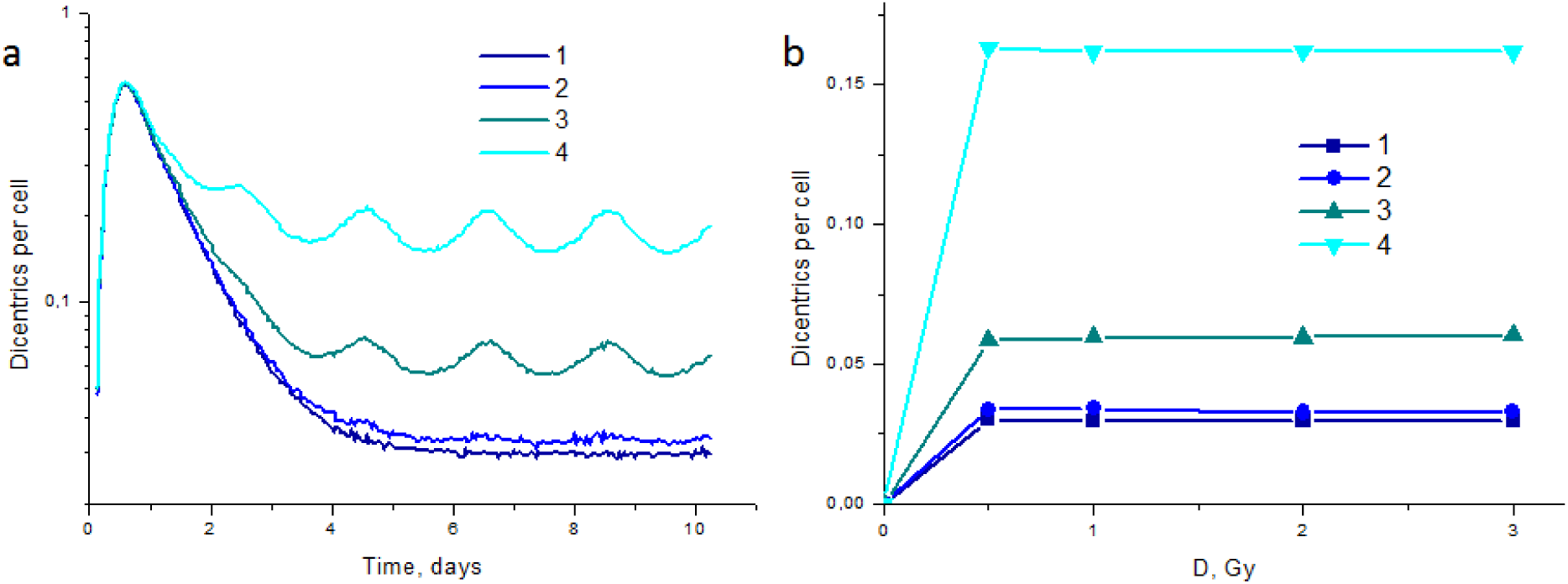
Non-autonomous DSB generation due to gap junctions) affects the dynamic curves of CI, but does not change the shape of the dose curves. (a): dynamic curves at a dose of 3 Gy for different contributions of autonomous and non-autonomous pathways of DSB generation. The autonomous level in all cases <n> = 5 per S phase. The level of non-autonomous DSB generation due to cellular contacts is varied. 1 - non-autonomous generation is absent (corresponds to curve 3 in Fig.1a). 2 - <n_GJ_> = 0.49 per S phase; 3 - <n_GJ_> = 4.9; 4 - <n_GJ_> = 24.5. (d2): dose dependencies at t = 10 days. The designations of curves are the same as in (a).

Deviations from the plateau form of the dose curves for a dose-independent DSB generation rate (Fig.1 g,h) occur when the plateau is not reached at large times by the dynamic curve, Fig.1 d,e. The reason for this behavior of the dose curve is investigated in Fig.5.

**Fig.5.**
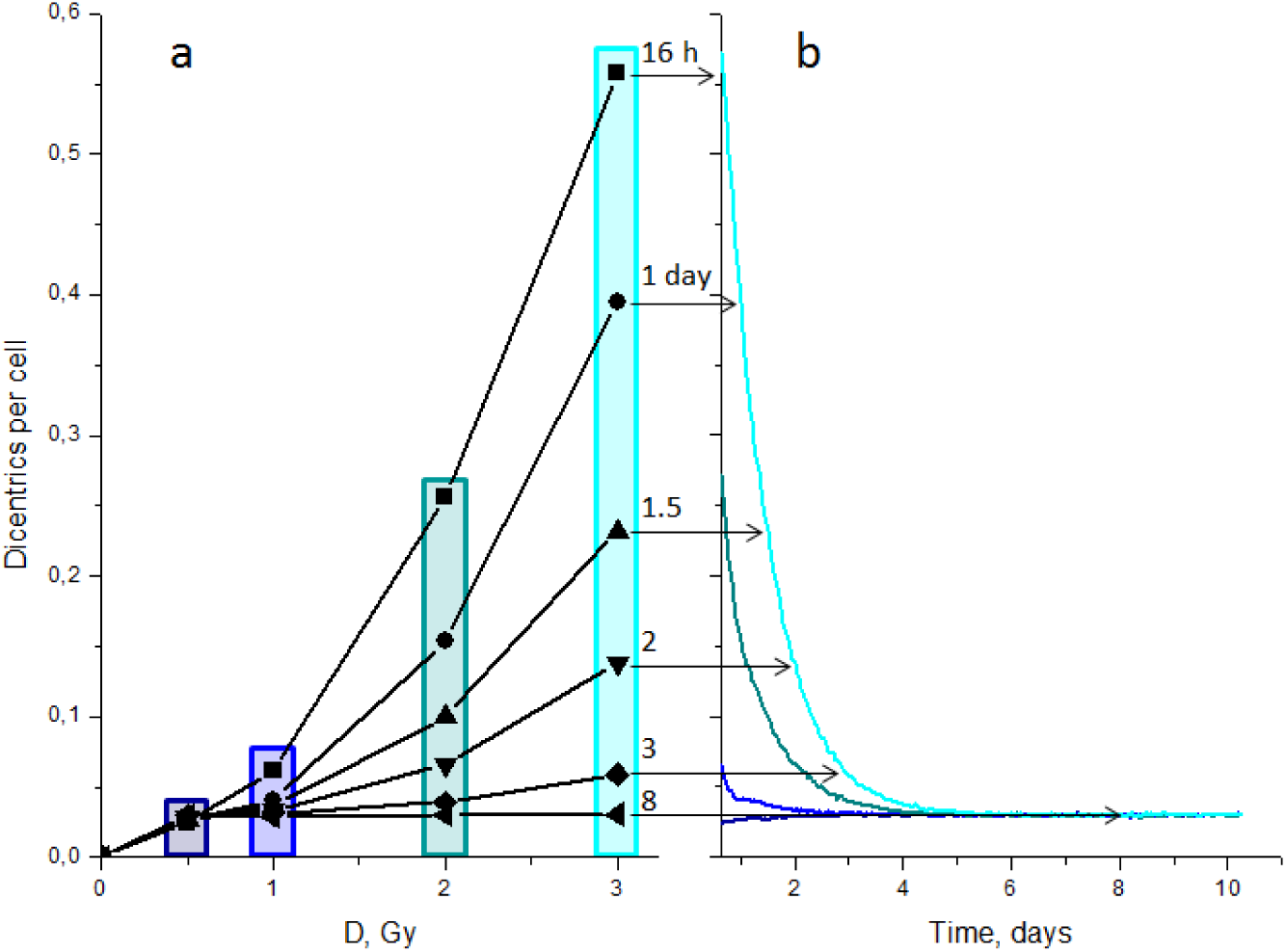
Dynamic origin of dose-response relationships for radiation induced CI, delayed dicentrics. (a): dose curves for different times post-irradiation. Colored rectangles highlight points for a single dose, but different times. (b): dynamic curves for different doses. The colors of the lines correspond to the highlight colors in (a). The arrows show the correspondence between two sets of curves for a dose of 3 Gy.

The theory shows how the dose dependence of CI is converted into dose independence. In the case of dose-independent rates of DSB generation and other parameters, the dose independence of CI is "converted" into dose dependence by a simple change in the observation time (the time of cell proliferation after irradiation), Fig.5.

This leads to an important prediction: if several factors/parameters change simultaneously, but the dynamics of dicentric generation-loss (dynamic curves) does not change for long times, the dose curve of CI remains unchanged. This prediction is verified by the calculations in Fig.6 for the third possible type of CI dynamics, similar to Fig.1d.

Two sets of dynamic curves, shown in Fig.6 a, are obtained for different sets of parameters (different values of three parameters). Coincidence of kinetic curves at all doses entails the coincidence of dose curves (Fig.6 e). To check whether we are in the region of system insensitivity to all three parameters, dynamic and dose curves were constructed for the cases when only one of these three parameters changes (Fig.6 b-d). It can be seen that the dynamic curves for all doses do not coincide, and this leads to a discrepancy between the dose curves at late times (Fig.6 f-h).

**Fig.6.**
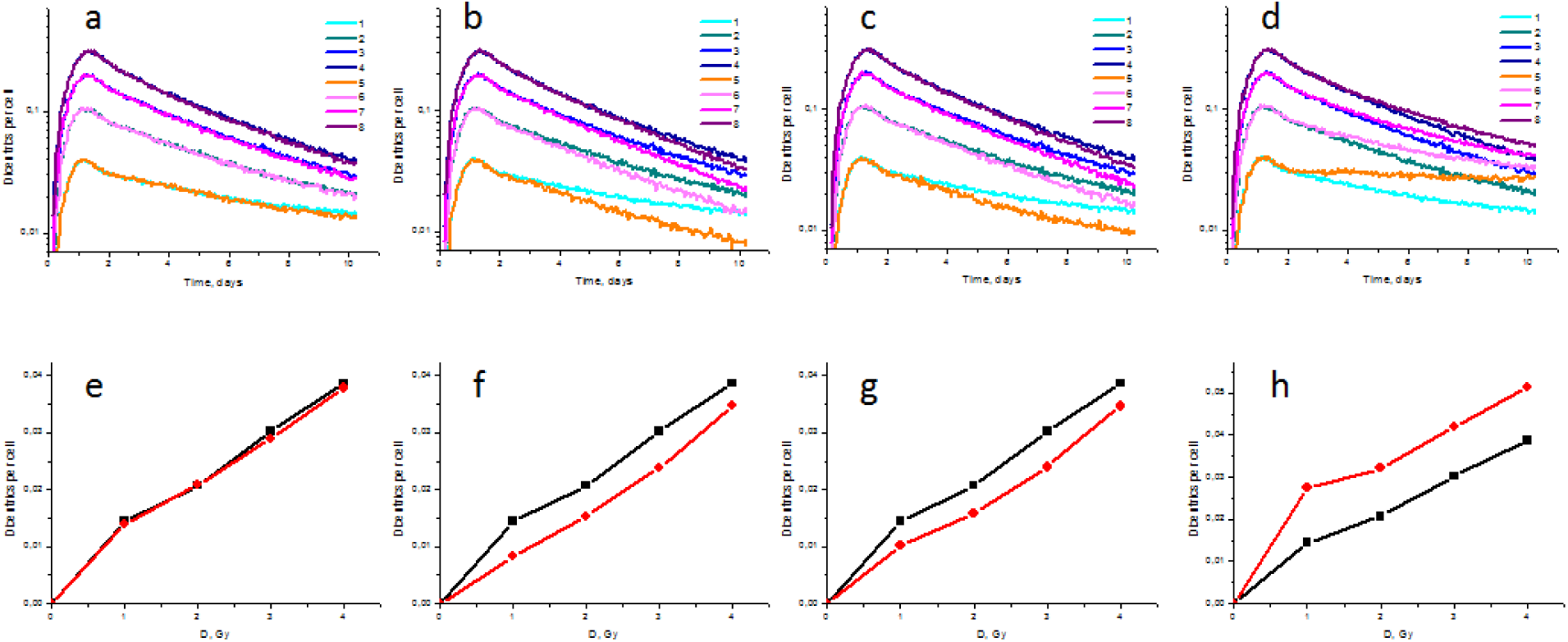
The dose dependence is directly determined by dynamic dependencies. (a) - (d): dynamic curves for doses 1-4 Gy obtained for different sets of parameters. (a): the three parameters change in such a way that the dynamic curves remain unchanged. The first set: generation of autonomous DSBs <n_auto_> = 2 per S phase; 100% of the DSBs are generated before the replicative fork; gap-junction-induced generation of non-autonomous DSBs <n_GJ_> is proportional to cell density, k = 0.18. The second set: <n_auto_> = 0.5; 20% of the DSBs are generated before the fork; k = 1.8. (b) - (d): only one of the three parameters changes. In all panels, the first set of parameters is the same as in the panel (a). The second set of parameters: (b) - changed <n_auto_>; (c) changed the fraction of DSBs before the fork; (d) - changed k. Doses in all panels: curves 1 and 5 - 1 Gy, curves 2 and 6 - 2 Gy, curves 3 and 7 - 3 Gy, curves 4 and 8 - 4 Gy. (e) - (h): dose curves at 10 days corresponding to the kinetic curves in (a) - (d). Black curves are obtained for the first set of parameters (see above); red curves, for the second set of parameters.

This demonstration of the deep relationship between the dose response and time dynamics was carried out for the dynamic CA curve which does not reach a plateau. The verification of this derivation for other types of dynamic curves will be carried out in separate studies dedicated to the analysis of experimental data.

In conclusion, the basic properties of the developed theoretical model of radiation induced CI can be formulated as follows:

1. Persistent induction of DNA DSBs and their repair determines the dynamic and dose characteristics of CI. They integrate both intra- and intercellular triggering factors of DSB generation and provide cellular pathways for their implementation in CI.
2. The dose curves of CI depend on the dynamics of accumulation and elimination of chromosomal aberrations. Thus, CI dose dependence-independence is of the dynamic nature. This is reflected in the fact that, for example, the dose dependence of the CI can be converted into dose independence (and *vice versa*), outside direct dependence on the contribution of intercellular signaling interactions and at dose-constant rates of all processes.
3. Changes in parameters affect the dose curves by affecting the dynamic curve, which is determined by the persistent formation and repair of DNA DSBs.

## Acknowledgements

The present work was supported by Russian Foundation for Basic Research grant 14-01-00825 to S.A.

S.A. acknowledges support from the MEPhI Academic Excellence Project (Contract No. 02.a03.21.0005).

## References

1 United Nations Scientific Committee on the Effects of Atomic Radiation (UNSCEAR). Effects of ionizing radiation. UNSCEAR 2006 Report.

2 M. Kadhim, S. Salomaa, E. Wright, G. Hildebrandt, O.V. Belyakov, K.M. Prise, M.P. Little. Non-targeted effects of ionising radiation - implications for low dose risk. Mutat. Res. 2013, V.752, pp.84–98.

3 S.G. Andreev, Y.A. Eidelman. Dose-response prediction for radiation-induced chromosomal instability. Radiat. Prot. Dosim. 2011, V.143, pp.270–273.

4 S.G. Andreev, Y.A. Eidelman, I.V. Salnikov, S.V. Slanina. Modeling Study of Dose-Response Relationships for Radiation-Induced Chromosomal Instability. Dokl. Biochem. Biophys. 2013, V.451, pp.171–175.

